# Premotor cortex hemodynamics reflect internal auditory category, not reported category

**DOI:** 10.1101/2024.12.18.628869

**Authors:** Jeff Boucher, Shihab Shamma, Yves Boubenec

## Abstract

In sensory decision-making tasks, animals’ decisions are driven by perception, but also by non-perceptual factors. Because of external and internal noise, stimuli may be internally misclassified, leading to perceptual errors. But other, non-sensory factors such as impulsivity or exploratory behavior can lead to non-perceptual errors. Here we exploited the neural traces of these errors in frontal cortex to provide insights into their role in sensory decision making. Using functional ultrasound imaging (fUS), we investigated how the premotor cortex (PMC) in ferrets represents stimuli in a categorization task, varying the difficulty in order to manipulate the rates of perceptual errors. We found that PMC activity reflects the objective (and not the chosen) stimulus category on incorrect Easy trials, when non-perceptual errors are more likely. In contrast, PMC responses correlate with the chosen category (and not objective category) on incorrect Difficult trials, when perceptual errors are more likely. These results suggest that PMC encodes the ferret’s perceptual decision but not necessarily the final motor decision. Perceptual errors could be refined further by assessing licking patterns, but licking patterns alone did not explain the effect. This study advances our understanding of the functional role of the frontal cortex in decision making, suggesting that the PMC integrates sensory inputs to guide behavior based on perceptual, rather than motivational, information.

## 2 Introduction

Perceptual decision making is the process by which sensory information is analyzed and converted into an action. In order to study this phenomenon, scientists often train their model animals on specifically designed tasks, with the intention that the animal reports their perception faithfully. A basic feature of such perceptual tasks is that the closer the perceptual objects to discriminate are, the more difficult the task and the more errors the participants will make [1]. These *stimulus-dependent* errors are often interpreted as evidence of the participants’ misperception, evidence that they have committed a *perceptual* error.

However, incorrect choices can also be due to *non-perceptual* (or *stimulus-independent*) errors. In humans, a common additional cause of such errors is a lapse in attention [2]. In addition to lapses of attention, a human may also intentionally make a wrong choice in order to explore options, rather than continuing to exploit a reliable pattern of behavior, if they expect that the optimal action might change [3]. Much more rarely (with the stronger effects in a neuropsychological context with lesions to the ventral PFC) [4–6], a human will demonstrate a strong bias for a previously rewarding action and be unable to prevent themselves from choosing it, even if sensory evidence obviously points in the other direction. In each of these cases, the action the participant took may not faithfully represent their true perception.

Non-human animals express each of these non-perceptual errors as well. Several studies have demonstrated that non-human animals show biases towards certain actions [7, 8] or explorative behavior [9]. In particular, in an appetitive Go/NoGo paradigm, an animal is expected to have a strong bias for the choice which grants them a reward [10], in a manner analogous to human patients. In these situations, the animals’ actions may not accurately report their perceptions.

In the ferret, the frontal cortex has recently begun to be investigated under auditory and visual perceptual tasks [11–14]. It has been shown that cells in both the prefrontal cortex (PFC) and the premotor cortex (PMC) respond in a categorical manner to task stimuli, responding to the stimulus’s meaning within the task rather than its physical parameters [12]. It remains an open question, however, whether the responses in this region tend to reflect the perception of the ferret or the final choice of action of the ferret.

Analysis of error trials might allow for a closing of this gap, particularly if we consider that not all error trials are equal: an error trial with a stimulus on the slope of the psychometric curve (a *Difficult* stimulus) should be relatively more likely to involve a *perceptual* error (Figure 1B); in this case, the stimulus would be internally classified as the incorrect category. For stimuli in the saturated regions of the psychometric curve (*Easy* stimuli), errors will relatively more often be *non-perceptual* errors rather than miscategorizations of the stimulus. As a consequence, it should be more probable in Easy error trials (relative to Difficult error trials) that a stimulus would internally be correctly categorized, despite the incorrect choice.

**Fig. 1.**
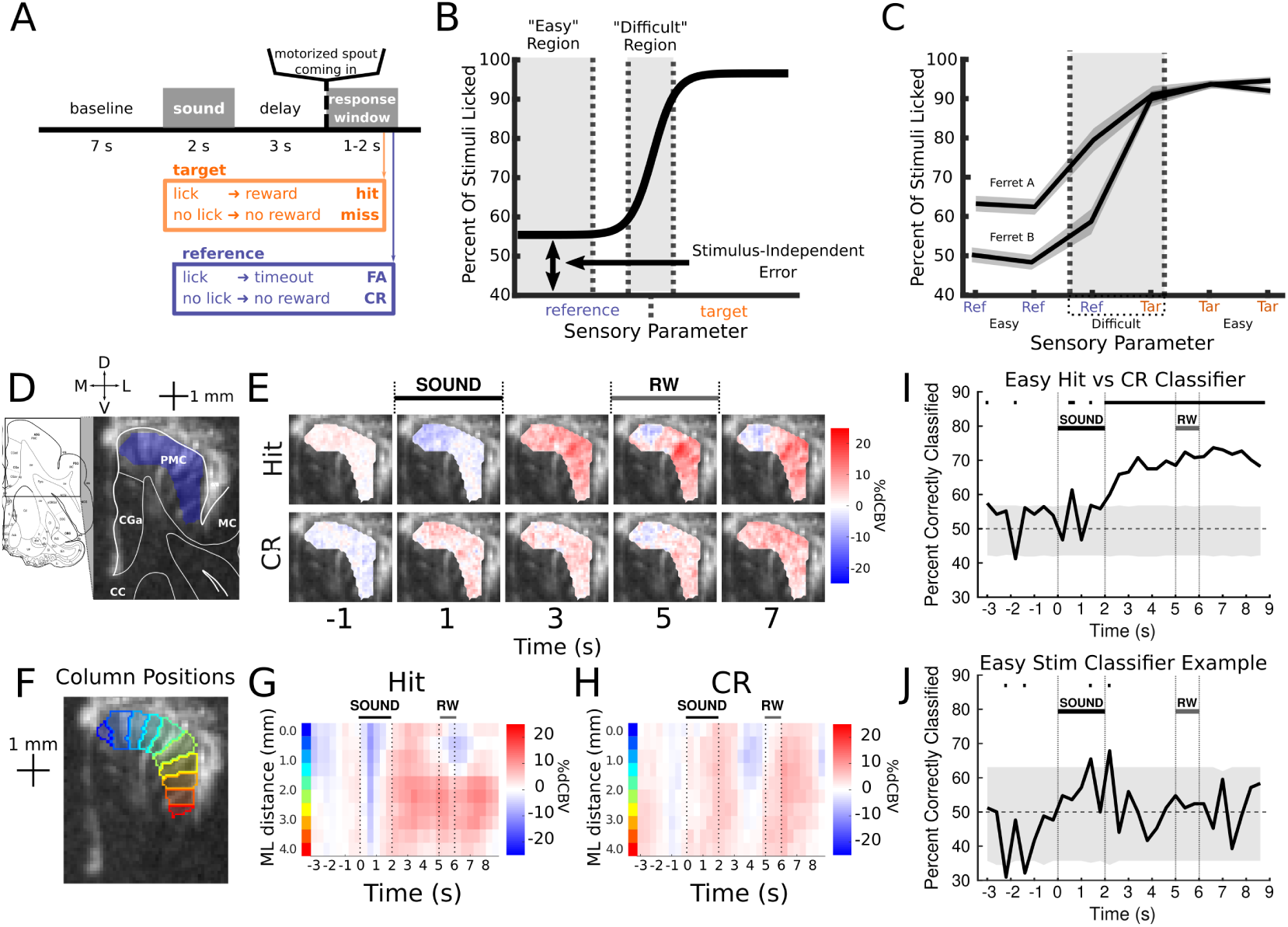
Task and Data Preparation. **A**: Cartoon describing trial structure. **B**: Cartoon illustrating the important parts of a psychometric function. For this binary Go/NoGo task, the two decisions depend on the value of the sensory parameter, and the correct decision can be defined at a specific boundary. At a far enough distance from this boundary (with the specific distance depending on the sensory parameter being studied) performance level becomes constant; errors may still be made, but they are independent of the stimulus and potentially independent of perception. Close to the boundary, errors increase; this is understood to be due to perceptions becoming more ambiguous in this region. **C**: Measured psychometric curves for ferrets A and B. **D**: Cartoon showing how to read the anatomy of a coronal fUS image. **E**: Example stimulus-evoked responses plotted per voxel. Voxel activity varies mainly along the surface of gray matter, and varies little across different depths. Only selected time bins are shown due to space concerns, but the full timecourse can be seen in panels G and H. **F**: Positions of the "columns" for the example session and slice. See Methods for how they were constructed. **G**: mean stimulus-evoked response for Hit trials, using the mean values of the columns in F. The important variations shown in panel E can be seen here, in addition to other time-frames. **H**: same as G, but for CR trials. **I**: The percent correctly classified for the Easy Hit-CR classifiers for each time frame. All sessions recorded at this slice were included to train the classifiers, including the example session of the prior figures. Classifiers were trained using the column means (visualized in panels G and H). The shaded area contains the 2.5% to 97.5% quantiles of the shuffled percents correctly classified. **J**: same as I, but for an example pair of two reference stimuli (specifically the two frequency-rule reference stimuli). All other pairs show similarly negative results, supporting the interpretation that responses are categorical.

Using this logic, we can disentangle the following: do PMC categorical responses reflect the category internally assigned to the stimulus, or do they reflect the enacted choice? If the choice is primarily what matters, all FA trials should show similar patterns of activity. If, however, the internal category of the stimulus is the primary driver of activity in PMC, activity during non-perceptual errors (more likely in FA on Easy trials) should look like that of the correct answers, and activity during perceptual errors (more likely in FA on Difficult trials) should look more like that of the opposite stimulus category (reflecting the miscategorization).

To address this question, we recorded hemodynamic responses in ferret frontal cortex during behavior with functional ultrasound imaging (fUS), a wide-scale hemodynamic imaging technique [15, 16]. Using fUS imaging, we scanned task-evoked neural responses in PMC while the ferrets performed a delayed-report Go/NoGo task (Fig 1). We were able to find distinct patterns of hemodynamic responses between correct Target (Hit) and Reference (Correct Rejections/CR) trials in PMC. Ferrets were trained to discriminate stimuli that were perceptually far from one another (Easy) as well as close to one another (Difficult). Easy and Difficult CR were similar, confirming that, when correct, Easy and Difficult trials showed categorical responses in PMC. Meanwhile, FA errors in Easy trials were also similar to CR trials, following the "stimulus category" hypothesis presented above. Finally, the Difficult FA trials were consistently closer to Hit trials than the Easy FA trials were, again in favor of the stimulus category hypothesis. This suggests that in ferret PMC, hemodynamic activity follows the pattern expected if it primarily responded to the internal sensory category, rather than the final decision made.

## 3 Materials and Methods

### 3.1 Animal preparation

For the purposes of this study, two ferrets (ferrets A and B) were implanted with stainless steel headposts fixed to their skulls, in a procedure standard to the laboratory’s prior work [17]. Experiments were approved by the French Ministry of Agriculture (protocol authorization: 21022) and strictly comply with the European directives on the protection of animals used for scientific purposes (2010/63/EU).

Following training, a frontal craniotomy was opened using a surgical micro-drill. Then, an ultrasound-transparent Polymethylpentene (TPXTM) plate was placed above the brain and underneath the skull, secured with dental cement. In order to maintain the position of the ultrasound probe relative to the ferret during behavior, a custom-built stainless steel (440C) ferromagnetic frame (Supp Fig 1) was cemented to the ferret’s implant surrounding the craniotomy, using a spirit level to keep it flat. The ultrasound probe could then be clamped to a paired magnetic frame, which could be magnetically attached to the ferret’s implant. This system also allowed for the use of a microtranslator for the repositioning of the probe along the anterior-posterior dimension while maintaining the rigid attachment to the ferret’s implant during their head-fixed behavior.

### 3.2 Stimuli and task

Both animals were trained on a Go/NoGo delayed response task. Each trial of the final task consisted of a 2-second Target or Reference Amplitude-Modulated (AM) pure tone, a 3-second post-stimulus silence, and the movement of the water spout in front of the ferret (Figure 1A). Licking the spout after a Target sound was rewarded with water. Licks after Reference sounds were punished with a 30 second time-out period before moving to the next trial. The valid lick-response window was 1 second for ferret A and 2 seconds for ferret B. An additional inter-trial interval of 7 seconds was also enforced, irrespective of the ferret’s choice.

Each ferret was trained to discriminate the AM tone stimuli along two different feature axes: carrier frequency or AM rate. During the "frequency rule" trials, all stimuli had a fixed AM rate of 5.8 Hz, and could have a carrier frequency selected from 521, 1094, 2514, 3520, 8089, or 17000 Hz. During the "AM rule", all stimuli had a fixed carrier frequency of 5657 Hz, and could have an AM rate selected from 1.2, 2.52, 4.8, 6.25, 11.9, or 25 Hz. Each ferret had to discriminate the three lowest AM rates or carrier frequencies from the three highest. Ferret A’s Target stimuli were low, while ferret B’s Target stimuli were high. Stimuli were chosen to be more than an octave apart from one another and to be easily discriminable across the category boundary, with the important exception of the two stimuli (per rule) close to the category boundary. These are called the "Difficult stimuli" as they were chosen to be more difficult to categorize. The remaining stimuli are called "Easy stimuli" due to the saturated performance levels. All stimulus positions were decided using the past behavioral results in [13].

Trials were organized into "feature-rule blocks". Within each block, ferrets could only be presented with sounds belonging to the prescribed rule. Blocks lasted 28 trials before switching; the 28 trials always consisted of 6 repetitions of each Easy stimulus and 2 repetitions of each Difficult stimulus, presented in pseudorandom order (with seed based on time). For every analysis, feature-rules are pooled together, as qualitatively the results within tasks were identical.

Ferrets were shaped on both rules from the first session, switching between them every 28 trials, with the Reference sound initially quieter than the Target sound and with there initially being no post-stimulus delay.

### 3.3 Behavior analyses

We used d’ to evaluate behavioral performance, a standard measure from signal detection theory [18, 19]. It is calculated by subtracting the inverse-norm of the False Alarm rate from the inverse-norm of the Hit rate. “Snooze” trials were excluded from all analyses. These were defined as sequences of consecutive trials for which the animal disengaged from the task. Because of the low criteria (as defined in signal detection theory) with which they performed the task (near-100% Hit rates), we used a highly conservative measure of snoozing, allowing for robust exclusion. We defined snooze blocks as any continuous sequence of trials for which the animals stopped licking that included at least 3 (missed) Go trials. If the ferret began to lick at stimuli again (Hit or FA), those trials and trials following them would be marked “non-snooze” unless the snooze criterion was again met. If the animal never licked until the end of the session, the last trial with a lick (Hit or FA) was the final non-snooze trial.

### 3.4 fUS analysis pre-preparation

#### 3.4.1 fUS recording and preprocessing

Ultrasound Images were collected using an Iconeus One system (Iconeus, Paris, France), using the default settings and the IcoPrime probe (15 MHz central frequency, 70% bandwidth, 0.110 mm pitch, 128 elements). The Iconeus One System collects images at 500 Hz and applies a spatio-temporal clutter filter [20] to isolate blood vessels from other tissues using sets of 200 contiguous images. After filtering, the machine takes the mean power over these 200 frames, outputting vascular images at an effective framerate of 2.5 Hz. All timebin labels refer to the beginning of this composite image; thus for example, the frame labeled "4.6 s" includes fUS data from 4.6 to 5.0 s.

#### 3.4.2 Correction for brain deformations using NoRMCorre

In fUS imaging, craniotomies are often quite large, and the brain is able to change shape along the course of a session. These movements tend to be slow, on the timescale of seconds. In order to correct for these movements, we applied the NoRMCorre algorithm [21, 22]. NoRMcorre is a “piecewise rigid template alignment” algorithm, designed to correct for movements which are inconsistent between distant regions of an image but consistent in the near-field, and specifically designed with two photon imaging in mind. The algorithm takes a template image and splits it into “patches”, each of which represents a smaller portion of the image. Every time-frame is then compared to the template patches and adjusted to the position of peak cross-correlation. These adjusted patches are then recombined through interpolation.

The algorithm was designed for use in correcting for movements of cells in two-photon imaging. Its reliability, therefore, is dependent on consistent landmark signals being present at each time frame and each patch. While large blood vessels can serve this purpose for fUS, they can be relatively sparse and are concentrated more visibly toward the surface of the brain. Patch sizes and overlaps were thus chosen conservatively, with a goal of correcting for vertical bulging movements of the brain. Specifically, every patch spanned the entire vertical range of each slice, and each was defined by a 600 micron central portion and 2400 microns of overlap on either side of that portion horizontally (medio-laterally). Patches were selected from within a cropped area of each fUS image containing all of the visible brain. These patches were adjusted vertically to counteract bulging movements that, when present, tended to be greater in the center of the craniotomy than toward the edge. Following the piecewise rigid template alignment, the patches were recombined using linear interpolation.

#### 3.4.3 ROI selection

The anatomical positions of each fUS recording were determined with the aid of a set of dedicated anatomical images, recorded on the first recording day following the implantation of the TPX. Coronal anatomical images of the entire craniotomy were taken with increments of 200 microns. The image depth was set to span the entire skull in order to aid in identifying landmarks. Each fUS recording position was determined by comparing the vasculature of example images from each session to the dedicated anatomical images and to histological slices reported in [23]. This process is illustrated in more detail in Supplementary Figure 2.

This study focuses on the PMC. The PMC ROIs were manually selected within each recording using the same histological images as references, and each session’s ROI was compared with a "template image" (one image per slice) to ensure similarity across sessions.

#### 3.4.4 Mean hemodynamic responses

All reported functional results are based on the measure “%dCBV” or “Percent change in cortical blood volume”. This is calculated for each time frame by subtracting the trial baseline mean of each voxel from the time frame, then dividing by the mean per voxel across all trials. The "baseline" time period was selected to be -1.8 seconds to -0.6 seconds inclusive relative to stimulus onset. This %dCBV measure is calculated after all preprocessing steps are completed, and is used to calculate the reported means per voxel.

#### 3.4.5 Vowel averaging within columns

This study additionally uses "columns" to organize the data for all major analyses (Figure 1F, G, H). Data collected with fUS shows strong correlations in responses over time between nearby pixels, in particular along the axis perpendicular to the gray matter surface. In order to better reflect this aspect of the data, increase signal to noise ratio, ease the pooling of data taken in different sessions, and decrease the number of variables inserted into later decoding models, voxels were placed into columns at fixed intervals along the cortical surface (Figure 1F), and the mean %dCBVs of each tile were analyzed directly. To do this, we used the "laynii" algorithm’s [24, 25] programs "LN_GROW_LAYERS" and "LN_COLUMNAR_DIST" to generate "columns" along the PMC after specifying the location of the surface and bottom of the ROI in a "rim" file and a seed column (on the medial edge of each ROI in this paper’s case). The average column size created in this manner was 0.125 mm.

In order to accurately match the columns across sessions for the classifier models, sets of four contiguous tiles would be combined into bins of size 0.5 mm, in a manner such that these larger bins would approximately align across sessions (Supp Fig 3). In order for this to work, an anchor vessel was selected in each slice (by visual prominence), and a "origin" tile was set to fully contain the anchor vessel in its center. As long as the initial ROIs were similar before tile generation, this was sufficient such that additional landmark vessels would be located in the proper bins on inspection (Supp Fig3). In order to verify this, all sessions were compared to a template set of bins for the session’s slice.

### 3.5 Classifier analyses

#### 3.5.1 Training and evaluation of classifiers

In order to assess similarities and differences in encoding of task variables in the PMC across conditions, we trained Linear Discriminant Analysis (LDA) models. All classifiers were trained to discriminate between "Hit" trials and "CR" trials, where the number of trials in each category was counterbalanced by randomly selecting a number of "Hit" trials equal to the "CR" trials (as CR trials were always fewer). For each frame and slice (but across sessions), the classifier was trained using MATLAB’s *fitcdiscr* function, with the columns’ %dCBV values as features. We evaluated each classifier using ten-fold cross-validation, resulting in a “percent test-set correctly classified” for each condition. These measures are evaluated against a permutation test: the labels of each trial are shuffled 10,000 times to create a nonparametric noise distribution. Direct statistical judgments can then be made by comparing the percent-correctly-classified against the (two-tailed) quantiles of the nonparametric distribution (Figure 1I). For all statistical tests performed on the later projection analyses, only frames for which the classification was above the 97.5th percentile were used. For the example stimulus-classifier plot, we underwent the same procedure for all pairs of Easy reference stimuli. The example plot shows the results from comparing the two frequency-rule tones at that slice; other slices and pairs showed similar results.

#### 3.5.2 Projection analyses

Analyses of projections always used trials (FA trials or Difficult CR trials) which were not directly involved in the training of the Hit/CR models. We projected the trials onto the average classifier across folds. For convenience, we plotted the timecourses of these projections in example plots (in the manner of Figure 2B, C, D) at all time frames in the trial, regardless of whether the classifier at that time-frame was significant. For the violin plots and statistical tests, we gathered summary statistics across time-frames (3.4-5.4 s post-stimulus window) and across slices. In order to do this, it was necessary to make the units of these classifiers comparable. To develop a method for this, the following reasoning was used: if a classifier is able to significantly distinguish between two categories, then the mean projected values of those two categories (Hit and CR in this case) may serve as reasonable anchors by which to align different axes to one another. To accomplish this, a scalar linear transformation was applied to each projected trial such that the mean projection of “Hit” trials would be 1 and the mean projection of “CR” trials would be -1. Specifically:

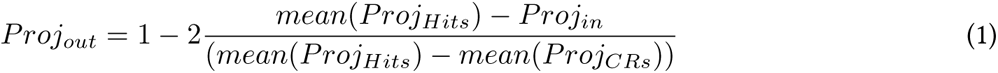

**Fig. 2.**
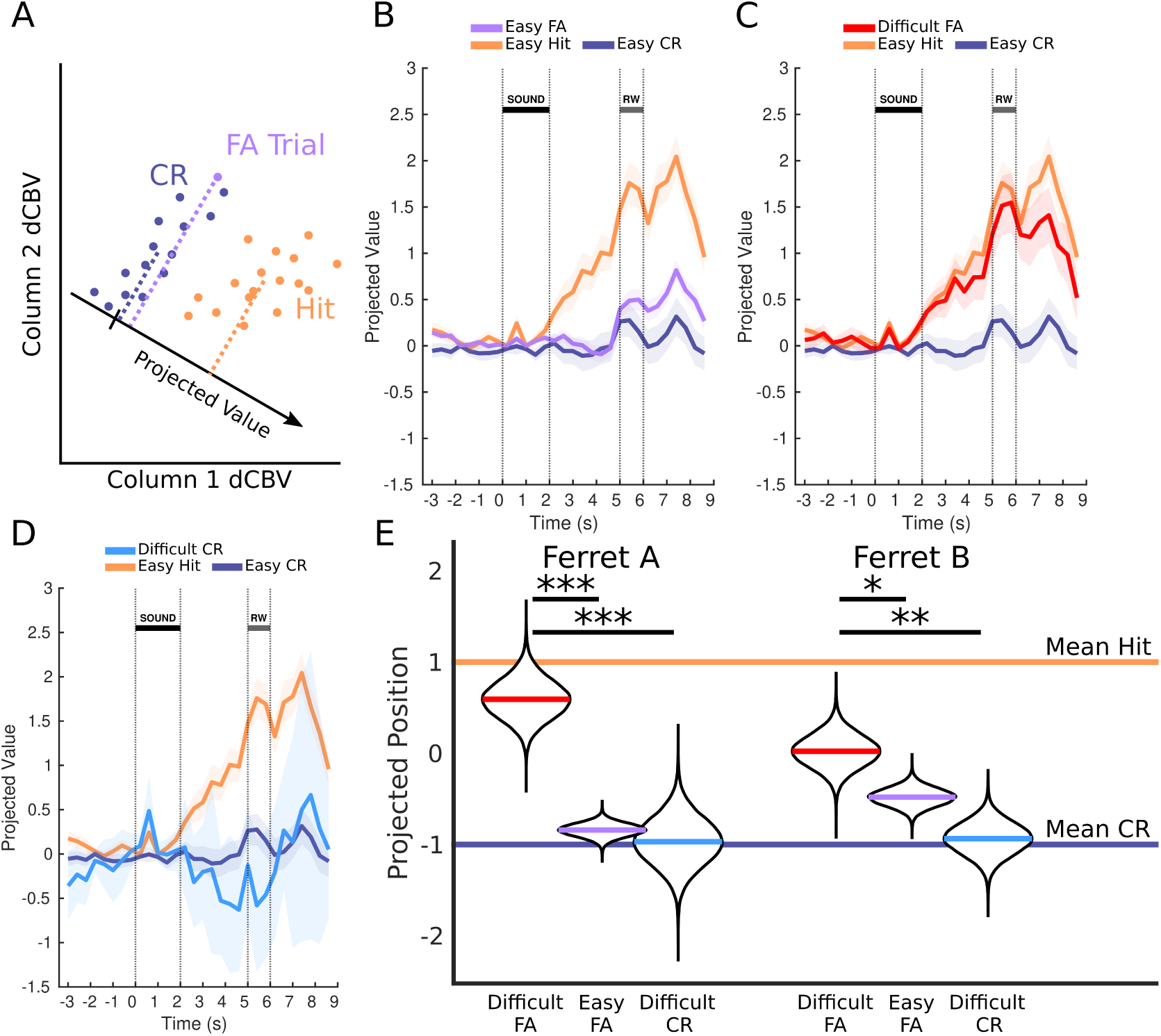
False Alarms project similarly to Correct Rejections for Easy but not for Difficult trials. **A**: Schematic of projections onto an optimal classification axis. The linear classifier will determine the direction in multiple dimensions (two are plotted here) for which two categories (Hit and CR) are classified. The positions of a Projected Value along this axis can then inform the extent to which the classifier would consider a trial to be a CR trial or a Hit trial. Arbitrary new trials, as long as they use comparable units (i.e., the same columns), can be meaningfully projected on this axis. **B**: Mean projections of PMC hemodynamic activity in Easy FA trials onto the example slice’s optimal axis. The mean projections of test-set Hit and CR trials are also plotted for reference. Shaded areas are 2 SEM. The HRF has an approximately 2-second delay, so interpretations of the timings should be adjusted to this. Note that the mean projection for Easy FA is aligned to Easy CR (same stimulus, different decisions), particularly in the seconds just following sound onset. **C**: Same as B, but for Difficult FA trials. Unlike Easy FA trials, Difficult FAs are aligned with the Easy Hit trials. **D**: Same as C, but for Difficult CR trials. The FA and CR projections for Difficult trials (C and D) do not look similar; Difficult CR trials look like Easy CR trials. **E**: Projections of the data aggregated across slices and across time frames from 3.4 to 5.4 seconds post-stimulus-onset (see Methods). All categories are colored the same as the prior panels; the mean Hit projection is enforced to be 1 and the mean CR is enforced to be -1 (see Methods), so they are plotted as horizontal lines. The bootstrapped means of Difficult FA trials for both ferrets are significantly separable from Easy FA trials and from Difficult CR trials.

**Fig. 3.**
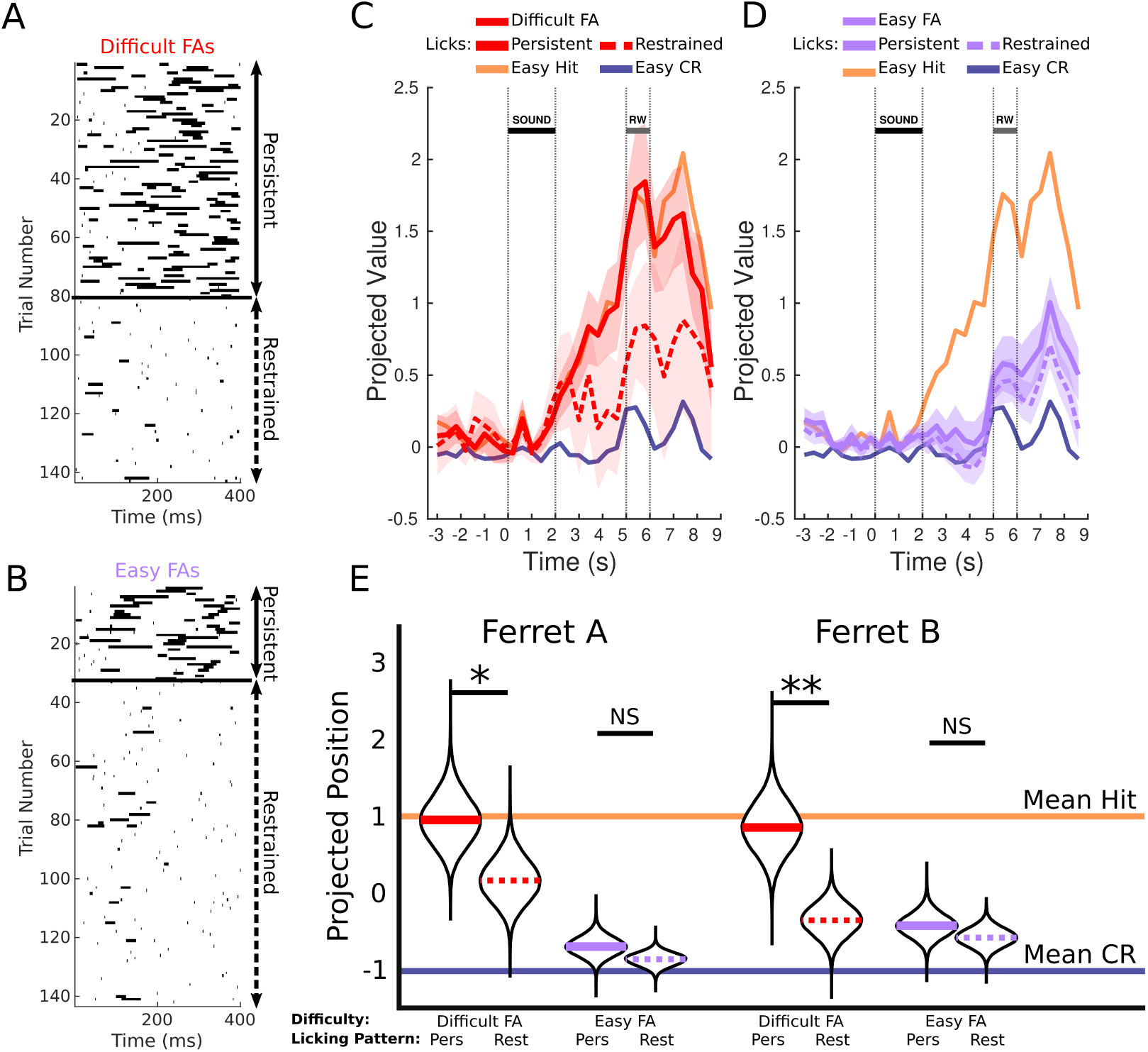
A characteristic licking behavior further refines perceptual errors in Difficult trials. **A**: Raster plot of licks for Difficult FA trials. Trials are sorted into "Persistent" and "Restrained". **B**: Same as A but for a randomly-selected subset of Easy-FA trials. **C**: Mean projections of PMC neural activity in Difficult FA trials (in the manner of Figure 2) grouped into "Persistent" licks and "Restrained" licks. When the ferret will restrain themselves following an initial FA in the reward window (dotted line), their Difficult FA activity projects closer to the Easy CR mean. When they make additional attempts, the projection is closer to the Hit mean. **D**: Same as C, but with Easy trials instead of Difficult trials. In the case of Easy trials, the "Persistent lick" behavior is less likely to reflect a misperception. Consistent with that idea, we see that Persistent licks and Restrained licks project to the same position for Easy FA trials, despite being distinguished using the same method as for Difficult trials. **E**: Projections of the data aggregated across slices and time frames (same format and method as Figure 2E). For both ferrets, Persistent lick trials project significantly closer to Hit trials than Restrained lick trials. For both ferrets, there is no distinguishing between Persistent and Restrained lick trials for the Easy FA trials.

Where Proj_Hits_ and Proj_CRs_ are vectors of the projections of all (test) trials for a frame’s classifier (Easy Hit and CR respectively) and Proj_in_ is whichever trial is currently being transformed (FA trials or Difficult CR trials). Projected trials with negative values would thus be closer to the mean CR projection, and projected trials with positive values would be closer to the mean Hit projection. It should again be stressed that this transformation can only be expected to be reasonable if the trained classifier is able to significantly distinguish the categories, otherwise the differences in mean projection for Hit and CR will be small and the remapping will be unwieldy. In order to test differences between the mean projected values of various conditions, we elected to use a hierarchical bootstrap procedure [26]. Specifically, we simulated 100,000 bootstrapped replications of the experiment by, for each chosen slice location, randomly pulling (with replacement) sessions and trials, taking the mean across all trials. The number of sessions randomly chosen per slice was equivalent to the number of recorded sessions per slice, and the number of trials per session was equivalent to the number of trials for the chosen session. The 100,000 means collected in this manner were used to create the distributions presented in the violin plot for each conditions. In order to test differences between means, we followed the procedure suggested in [26]: we randomly paired bootstrapped means from each distribution to create a joint distribution, and tested to see the percentage of samples were on the appropriate side of the unit line (all tests were one sided). Asterisks reflect one-sided tests: "*" means less than 4.55%, "**" means less than 0.27%, and "***" means less than 0.0063% (2, 3, and 4 standard deviations).

#### 3.5.3 Regression control for the influence of motor activity

The ridge regression models used for the regression control were built from five variables: lick rate at four delays (1.2, 1.6, 2.0, and 2.4 seconds preceding the time frame) and motion energy at the moment of the recording frame. The ridge regression code used was *ridgeMML* from [27]. The manner in which lick rate was calculated will be discussed in the section "Video analysis of licking" below. Motion energy was calculated by taking the absolute difference between consecutive frames per pixel, and taking the mean across all pixels. Ridge regression models were built independently for each column and each frame, but across sessions for each column. All non-snooze trials were used to construct the models, agnostic of difficulty or response.

After building these models, we used them to predict the values of each column by first multiplying the regression weights by the regressor values for each trial and column (after subtracting the mean of each regressor across trials). This reconstructs the mean column values recentered around the mean; therefore we re-added the means to get the final reconstructed values. These full predicted models were then subtracted from the actual recorded data, removing the components which could be explained by the ridge regression model. All classifier construction and projection steps were then repeated with the residual data.

### 3.6 Lick and motion analyses

#### 3.6.1 Video analysis of licking

In order to evaluate the potential contributions of lick evoked or lick preparatory activity, lick rates had to be collected in the period before the spout arrived. To accomplish this, video data was recorded at a framerate of 7.5 Hz. Each frame was aligned to the fUS data by using an LED set to light up each second starting at the beginning of each trial and ending before the inter-trial interval (ITI). Lick rate was calculated by drawing ROIs around the mouth of each ferret in each video (Figure 4A) and looking for changes in the color pink from frame to frame (in order to simply but effectively filter out whisker movements). This color difference was calculated by subtracting the blue signal from the red signal for each frame, then subtracting consecutive frames from one another and taking the absolute value. The threshold for making the lick judgments was set by eye per video, in order to give the best match to actual licks for the first ten trials of each session when watching a video of the ferret. Each video frame was evaluated as having or not having a lick in a binary manner; these assessments were summed in order to downsample into the frame period of the fUS acquisition (33 ms into 600 ms or 133 ms into 400 ms). For the violin plots in Figure 4B, we took the mean of this lick rate across frames 1-4.2 seconds for every trial and for each analyzed stimulus/response type.

**Fig. 4.**
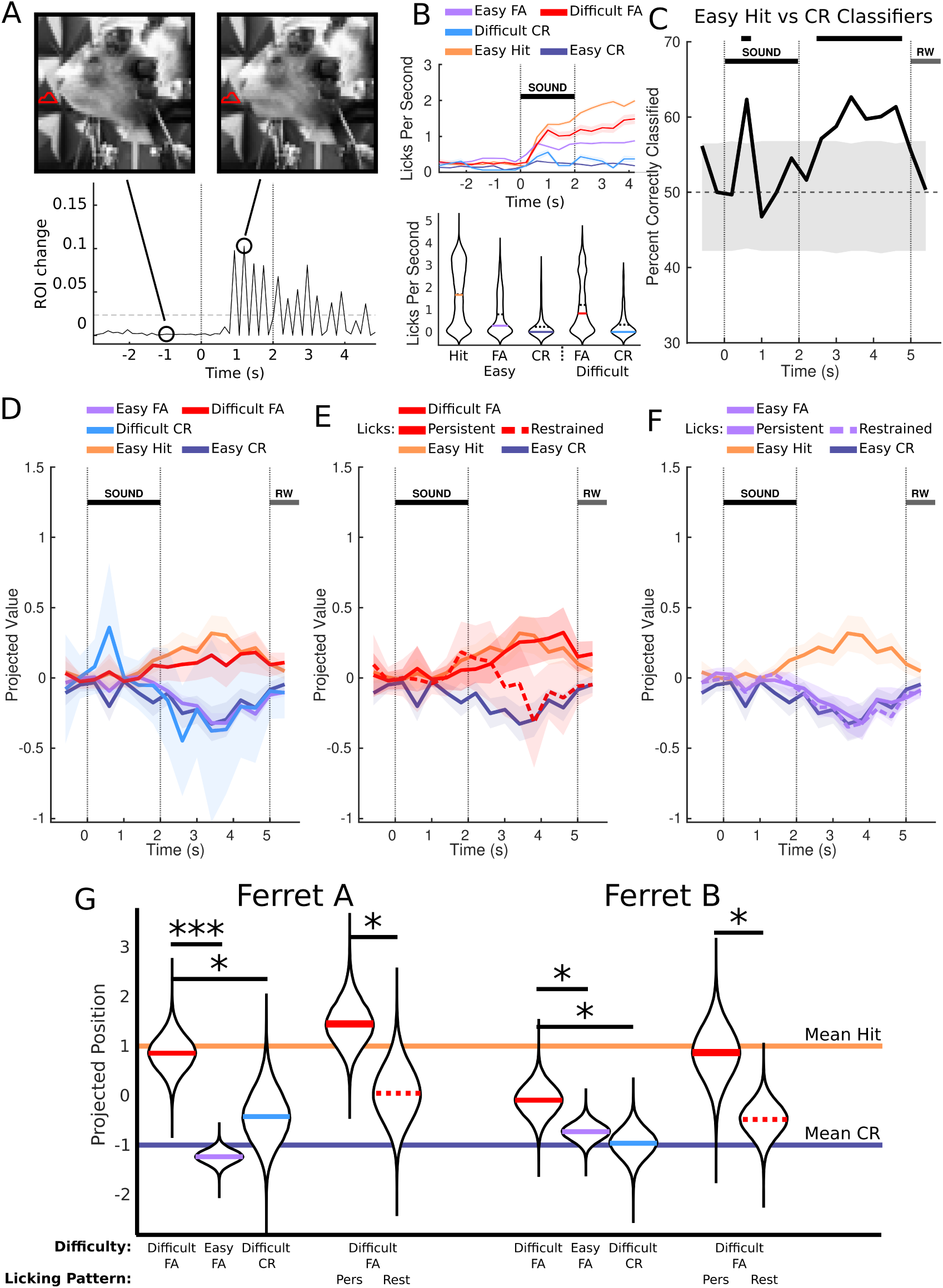
Pre-spout licking cannot explain the difficulty effects. **A**: Illustration of the video lick detection method. We drew ROIs where licks might occur, and tracked changes in them (see Methods). **B**: Top: Average lick rates for Easy Hits, Easy CRs, and Easy FAs (ferret A). Shaded areas are the standard error. Bottom: Violin plots showing average lick rate in each trial over the period from 1 to 4.6 seconds. The dotted line is the mean, the colored is the median. **C**: Easy Hit-CR decoding accuracy from PMC neural activity (as in Figure 1I) after subtracting the movement-related contribution. The absolute magnitude of the classification has been reduced, suggesting that lick rate contributed to the classification ability. However, classification is still significant in the post-stimulus period, suggesting differentiating activity unrelated to this. **D**: Mean projections of PMC neural activity for Difficult FA, Easy FA, and Difficult CR (as in Figure 2B, C, and D) using the regression-subtraction classifiers. Difficult FA trials still project close to the mean Hit projections, and Easy FA and Difficult CR trials still project to the mean Easy CR projection. **E**: Persistent/Restrained Lick trial projection plot for Difficult FA trials (as in Figure 3A). Restrained licks still project close to Easy CR trials, despite being Difficult FA trials. **F**: Persistent/Restrained Lick trial projection plot for Easy FA trials (as in Figure 3B). The licking pattern on spout arrival does not affect the projected value of Easy FA trials. **G**: Projections of the data aggregated across slices. Persistent/Restrained Lick trials for Easy FA are omitted for space, but would be aligned with the Easy FA means. All results previously reported are maintained.

#### 3.6.2 Determination of Persistent and Restrained licking

Persistent and Restrained licking were defined in terms of licking behavior following the spout arrival. Licking intensity following spout arrival was quantified using a piezoelectric actuator attached to the water spout, which detected the mechanical vibrations caused by licks. Conceptually, Persistent licking was intended to capture continuous licking after an initial lick, either in a ferret’s attempt to retrigger the piezo for an expected Hit, or because the ferret hoped to catch expected water. Persistent licking was thus defined as any trial where the piezoelectric device was activated for more than 30 ms in the period 200-400 ms following the response window, while all other FA trials were categorized as Restrained licking trials. This definition would also include other licking styles that extended into the later response window, such as a continuous licking unconcerned with the stimulus that may occur at the beginning of a session, but it was believed that in the case of perceptual errors the intended licking styles would occur often enough for the metric to capture them.

The threshold of 30 ms was chosen to capture all trials where one or multiple prolonged licks occurred in the 200-400 ms response window. Due to the piezoelectric device’s inherent mechanical ringing properties, it exhibited rapid and intermittent on/off patterns even following single licks. To accurately capture periods of continuous licking, piezo activations occurring within a 37 ms interval were considered as part of a single lick bout and time frames in between were flagged as "activated".

## 4 Results

### 4.1 Ferrets perform tasks which spans the psychometric curve

Ferrets were trained on a Go/No-Go delayed categorization task under appetitive reinforcement. Water-deprived ferrets had to classify 2 second-long AM pure tones into two categories: “Target” (signaling the ferret to lick) or “Reference” (signaling the ferret to restrain themselves). Stimuli were followed by a 3 second delay during which the water spout was out of reach (Figure 1A). Licks during the subsequent response window were rewarded with water in Target Hit trials and punished with a timeout in Reference False Alarm (FA) trials. The task was designed to measure the ferrets’ response accuracy through a classic sigmoid-shaped psychometric curve, reflecting how error rates vary with stimulus parameters (Figure 1B). Performance was saturated for the “Easy” Target stimuli (ferret A: d’ = 1.2 with Hit rate 0.94 *±* 0.007, n = 1120 trials, FA rate 0.63 *±* 0.014, n = 1182 trials; ferret B: d’ = 1.48 with Hit rate 0.93 *±* 0.007, n = 1353 trials, FA rate 0.49 *±* 0.013, n = 1409 trials; Figure 1C). Meanwhile the performance for the “Difficult” Reference stimuli, placed closer to the category boundary, was worse compared to the Easy Reference stimuli (Figure 1C). Trials for which the ferrets ceased licking were excluded (snooze trials, see Methods), but blocks of trials or sessions during which the ferrets had high FA rates were included, allowing for detailed analysis of trials where False Alarms were made on Easy stimuli. Both ferrets had near-100% Hit rates, suggesting low decision criteria; due to this, analysis of error trials was focused on Reference False Alarms (FAs), rather than Misses. Ferrets made significantly more FAs on the Difficult Reference stimuli compared with the Easy Reference stimuli (ferret A p < 10^-5^, n = 193 Difficult trials, 1182 Easy trials; ferret B p = 0.004, n = 235 Difficult trials, 1409 Easy trials, one-tailed two sample z-test), confirming that the ferrets found the stimuli more difficult to classify. In contrast, FA rates were indistinguishable between the Easy Reference stimuli (ferret A p = 0.78, n = 590 Easy ref stim 1, 592 Easy ref stim 2; ferret B p = 0.49, n = 711 Easy stim 1, 698 Easy stim 2, two-tailed two sample z-test), consistent with saturated performance.

### 4.2 Large-scale ferret premotor cortex hemodynamics distinguish between Hit and Correct Rejection trials during active behavior

We first established differences in the large-scale hemodynamic responses during correct Target (Hit) and Reference (Correct Rejection; CR) trials. We recorded changes in blood volume (%dCBV) in premotor cortex (PMC) of the two behaving ferrets using fUS neuroimaging (Figure 1D,E). Recordings were performed repeatedly at specific positions in PMC (34 sessions at 2 positions for ferret A; 37 sessions in 3 positions for ferret B), selected following a preliminary study. These positions could be mapped to the ferret brain atlas [23] through anatomical landmarks (Supp Figure 2). For the remainder of the paper, one position of ferret A (n = 17 sessions) will be used for all examples, though the results generalize between slices and ferrets (provided in the supplementary figures).

We found distinct hemodynamic responses in PMC between Hit and CR trials. Target sounds evoked a transient suppression at around 1 s (Figure 1E) followed by a sustained increase in CBV persisting throughout the delay. This pattern remained clear after pooling voxels into 0.5 mm-wide columns defined by local anatomy, allowing data to be aligned by these columns across sessions while maintaining the Hit-CR difference (Figure 1F-H; see Methods and Supp Figure 3). Interestingly, we noticed some spatial heterogeneity across the mediolateral axis, in particular after the stimulus period (Figure 1E,G). We therefore used a decoding approach to examine the neural representation of Hit and CR trials. To do so, we performed linear decoding to determine time-dependent optimal linear classification axes between Hit and CR trials (Figure 1I). Decoding accuracy between Hit and CR became significant 2 seconds after stimulus onset and increased to a plateau about 1 second later. As the hemodynamic response function (HRF) for cortical blood volume (CBV) is expected to peak at about 2 seconds after the onset of neural responses [17, 28], this suggests that the CBV-based classifiers’ ability to discriminate between Hit and CR are consistent with the latencies of the stimulus-evoked responses at the single-neuron level in previous PMC studies [12, 13]. Furthermore, stimuli from within the same category could not be decoded from each other (Figure 1J), confirming that PMC responses are mostly categorical.

### 4.3 False Alarms project similarly to Correct Rejections for Easy but not for Difficult trials

We investigated whether ferret PMC activity correlated more with the internally-classified categories or with expressed behavior by examining False Alarms during Easy and Difficult trials. False Alarms made in the steep region of the psychometric function (Difficult trials) are more likely to be due to genuine perceptual miscategorization compared with those made in the saturated region (Easy trials). However, False Alarms in either case involve a lick action. If the evoked activity during Difficult and Easy False Alarms were similar, this may imply that the decision to lick was driving the hemodynamic response. We hypothesized, however, that PMC activity would primarily reflect the ferret’s internally classified category for the stimulus. If this were the case, we would expect PMC categorical responses to represent the *Target*category in a Difficult *Reference* FA trial, as *perceptual* errors are more likely. However, in Easy *Reference* FA trials, PMC activity would look more similar to the *Reference* stimulus, as *non-perceptual* errors should be more likely.

To test this hypothesis, we assessed whether Easy or Difficult FA trials were more similar to Hit or CR trials by projecting them on the Hit-CR classifiers trained for each slice and time point across sessions (Figure 1I, Figure 2). This procedure reduces the multicolumn hemodynamic PMC responses to a single dimension encoding the similarity to Hit or CRs patterns (Figure 2A). In addition to projecting the FA trials, we additionally projected the test-set Hit and CR trials of each fold onto the fold’s axis for illustrative purposes. Easy FA trials show activity more similar to that of CR trials than to Hit trials despite Hit trials sharing the decision to lick (Figure 2B, Supp Figures 5B, F, J, N). In contrast, in Difficult FA trials, where a misperception of the stimulus category is more likely, projections are more similar to Hits (Figure 2C, Supp Figures 5B, F, J, N). This is not likely to be an effect of stimulus identity, however, because Difficult and Easy reference trials align for Correct Rejections (Figure 2D, Supp Figures 5B, F, J, N).

The mean projection of Difficult FA trials is significantly closer to the mean Hit projection than Easy FA trials for both ferrets and across recording positions (Figure 2E; ferret A p < 10^-5^, ferret B p = 0.007). Additionally, the mean projection of Difficult FA trials is significantly closer to the mean Hit projection than Difficult CR trials (ferret A p < 10^-5^, ferret B p < 10^-4^). To allow for data aggregation across sessions, we normalized each slice by adjusting the projections so the mean "Hit" was 1 and the mean "CR" (Correct Rejection) was -1 (Figure 2E). This normalization applied uniformly to all trials, ensuring comparisons between trials reflected their proximity to these normalized values without altering their distribution. We then calculated bootstrapped means (see Methods) for the evaluated trial types—Difficult FA, Easy FA, and Difficult CR—to assess the significance of observed effects across slices, focusing on the transition into the delay period (3.4-5.4 s post-stimulus onset). Altogether, these results argue against the idea that the activity primarily reflects the expressed decision, and argue in favor of the idea that the activity reflects the internal classification of the stimulus category.

### 4.4 A characteristic licking behavior further refines perceptual errors in Difficult trials

So far, we have introduced the concept that False Alarms can arise from multiple causes, and that many False Alarms in Difficult trials can stem from perceptual miscategorization. To further refine our analysis of such trials, we examined licking behaviors, which potentially indicate the animal’s confidence in its decision and expectation of a reward. We observed that during False Alarm trials, ferrets often engaged in "Persistent" licking (Figure 3A, B and Supp Figures 4), continuously licking the spout despite the lack of water, suggesting either an anticipation of receiving a water reward or a bias toward licking. Conversely, at times, the animals exhibited "Restrained" licks (Figure 3A, B and Supp Figures 4), where they licked the spout just once. We hypothesized that Persistent licks in Difficult trials are more likely to be associated with perceptual miscategorizations than Restrained licks. Consequently, these distinct licking patterns may facilitate the differentiation between perceptual and non-perceptual errors in Difficult trials.

We split FA trials into "Persistent Lick" trials and "Restrained Lick" trials depending on whether extended licking behavior was observed in the lick detector during the late response window (see Methods). We then separately projected the split Difficult FA trials onto the same classification axes as in the prior analysis (Figure 3C, Supp Figures 5C, G, K, O). We found that the Persistent Lick trials projected closer to the Hit projections compared to Restrained Lick trials for all slices and ferrets (Figure 3E, leftmost red-colored violin plots; ferret A p = 0.033, ferret B p = 0.0008).

By design, Persistent licking trials involved more pronounced motor activity than the Restrained licking trials. We wanted to control for the possibility that the refinement of Difficult trials into Hit-and CR-aligned groups based on licking behavior was not merely dependent on differential motor activity. Given that the stimuli in Easy trials lack perceptual ambiguity, we hypothesized that Persistent licks in these Easy False Alarm trials were predominantly influenced by biased licking behaviors rather than perceptual errors. Consequently, we predicted that the majority of False Alarms in Easy trials, regardless of licking type, would represent non-perceptual errors.

To test this hypothesis, we categorized Easy FA trials into "Persistent Lick" and "Restrained Lick" trials in the same manner, projecting these onto the Hit/CR classifiers (Figure 3D). Contrary to Difficult trials, no significant difference was observed in the mean projection values between the two types of Easy trials (Figure 3E, rightmost lavender-colored plots; ferret A p = 0.14, ferret B p = 0.21). This analysis supports the notion that licking behavior specifically refines perceptual and non-perceptual errors in Difficult FA trials.

### 4.5 Lick-related activity cannot explain the difficulty effect

Ferrets were not punished or rewarded for licking outside of the response window. They sometimes chose to lick in response to the stimulus before the spout arrived (Figure 4A). This behavior was dependent on the stimulus and on the ferret’s future response in each trial (Figure 4B, top), though there was a large variability between trials (Figure 4B, bottom). In order to account for the possibility that this may affect the classification axes, we proceeded to remove all lick-related activity using ridge regression models. These models included both (i) regressors created from the ferrets’ lick rates at multiple delays (1.2, 1.6, 2.0, and 2.4 seconds preceding the time frame), accounting for potential lick-triggered hemodynamic responses, and (ii) a regressor created from the motion energy of the video (zero-lag), accounting for any remaining motion artifacts (see Methods). We then subtracted the contribution of these regressors from the hemodynamic activity and reran the full analysis, retraining the Hit-CR classifiers and projecting the False Alarm data onto them. The classifiers were still able to significantly classify Hit vs CR without the subtracted components in the relevant time period of 3.4-5.4 seconds for all ferrets and slices (Figure 4C, Supp Figures 6A, E, I, M), excepting a few frames post-spout-arrival for two slices (Figure 4C, Supp Figure 6M), allowing us to compare the original and subtracted neural data.

Qualitatively, the results are all consistent with those obtained before regressing out lick-related activity (Figures 2 and 3). All projection analyses presented thus far are provided in the regression-subtraction context (Figures 4D-G, Supp Figure 6). The main result that Difficult FA activity projects closer to the mean Hit activity than Easy FA trials (ferret A p < 10^-5^, ferret B p = 0.0428) or Difficult CR trials (ferret A p = 0.0256, ferret B p = 0.0207) is robust. The Persistent licking result that, among Difficult FA trials, Persistent Lick trials project closer to the mean Hit projection than Restrained Lick trials (ferret A p = 0.0230, ferret B p = 0.0166) is also maintained. Finally, the result that, among Easy FA trials, there is no significant difference between Persistent Lick trials and Restrained Lick trials (ferret A p = 0.2119, ferret B p = 0.3424) is also maintained. For these reasons, we believe that, while the licking behavior may have contributed somewhat to the shape of the classification axis, the placement of perceptual and non-perceptual errors along optimal classification axes are unaffected.

## 5 Discussion

Perceptual decision making, in addition to the perceptual factors most often studied, involves many non-perceptual factors. While disentangling these can be difficult on a per-trial basis, established theoretical models of behavior using signal detection theory allow us to have experimental control of the likelihood of errors caused by each kind of factor. In this study, we were able to leverage this differential proportion of perceptual errors to demonstrate that the hemodynamic response profiles of the ferret PMC primarily reflect the perceptual factors of decision making, instead of the animal’s motor report. We did this by demonstrating that the hemodynamic response during False Alarm trials depended on difficulty, which is highly correlated with the likelihood of perceptual errors.

### 5.1 fUS imaging of PMC in behaving ferrets

We used functional ultrasound imaging (fUS) to explore how the PMC encodes sensory evidence and categorization during decision making. This confirms the possibility of this recent imaging modality in behaving animals, that has been used only in the primate to our knowledge [29]. Notably, fUS imaging, while offering high spatial and temporal resolution compared to other hemodynamic-based imaging techniques, measures neural activity indirectly through blood flow changes rather than direct neuronal activity. This hemodynamic response is generally believed to reflect the aggregate activity of large neural populations rather than the activity of individual neurons. Consequently, our findings reveal the integrated response of the PMC to sensory stimuli, capturing a broader neural activation. This methodological approach provides insights into the overall brain state changes and metabolic demands associated with different cognitive processes, but it might mask the finer, more rapid fluctuations in neural activity that direct neuronal recording techniques like electrophysiology could detect. Therefore, while we observe that the PMC accurately encodes perceptual categories even during incorrect decision-making, these results must be interpreted with an understanding that the hemodynamic signals might not capture all aspects of the underlying neural dynamics, particularly those driven by fast, transient neuronal interactions that could be crucial for decision-making processes.

### 5.2 Categorization in PMC and fluctuations in internal states

Frontal regions are known to be involved in response inhibition through the hyperdirect pathway that projects to the subthalamic nucleus [30, 31]. Surprisingly, we found that even when the PMC accurately encodes the correct sensory category on No-Go trials, ferrets may still make incorrect decisions. It suggests that the PMC signal to suppress responses to No-Go sounds might be shortcut on FA trials. This observation is consistent with the results of [32], who reported that decision-making based on sensory evidence is not necessarily contaminated by internal state fluctuations such as impulsivity. In their study, neural activity drift observed in the visual and prefrontal cortices acted as a brain-wide impulsivity signal that, notably, would not alter the encoding of sensory stimuli but rather would modulate decision making independently. This suggests a mechanism where sensory perception within the PMC remains intact and unbiased by internal states, consistent with our findings where the PMC represented sensory evidence correctly even when ferrets made incorrect choices. We hypothesize that while the PMC provides an accurate sensory representation, decision errors may arise from downstream circuits that are intermittently overridden by internal states like impulsivity. These internal states may influence the decision-making process independently of the sensory evidence, leading to perceptual errors.

### 5.3 Nature and neural origin of non-perceptual decision making

In the prior section we proposed that downstream circuits may override the categorical signal in PMC in order to result in a non-perceptual error. Where might these downstream circuits be? This likely depends on the sort of lick decision that is made. Typically, parietal regions are thought to be involved in driving history-based exploratory behaviors [33–35], and so these areas may contribute to overriding the PMC signal for some trials. In addition, the "restrained Licks" as defined in this study seem to be more of a reflex than a deliberation: they involve just one lick, timed to spout arrival, followed by a quick restraint. An area involved with such a reflexive action may perhaps have a close if not direct connection to medial medullary RF (an area shown to be the central pattern generator for licking in cats [36]) in order to manage this speed and override the slower processes. A potential region which meets this criterion could be orbital frontal cortex. This region, being agranular frontal cortex, is highly connected to the limbic system, making plausible an involvement in managing criterions for rewarding stimuli; furthermore, in the cat, microstimulating a frontal orbital cortical region resulted in rhythmic licking behavior [37]. Unfortunately, these areas were beyond our imaging window. Future studies should focus on these regions to elucidate the comparative roles of parietal and fronto-orbital regions in decision making across rodents and carnivores.

## Acknowledgements

We would like to thank Thibaut Doisy, Denis Lancelin, Gwenola Dubois, Balkis Cadi, and Lynda Bourguignon for technical and administrative support throughout the project. This work was supported by ANR-17-EURE-0017 and ANR-10-IDEX-0001-02, CDSN doctoral fellowship from École Normale Supérieure to JB, ERC 787836-NEUME and ANR-19-CE37-0016 to SAS, and ANR-JCJC-DynaMiC to YB.

## Supplementary information

**Supplementary Figure 1.**
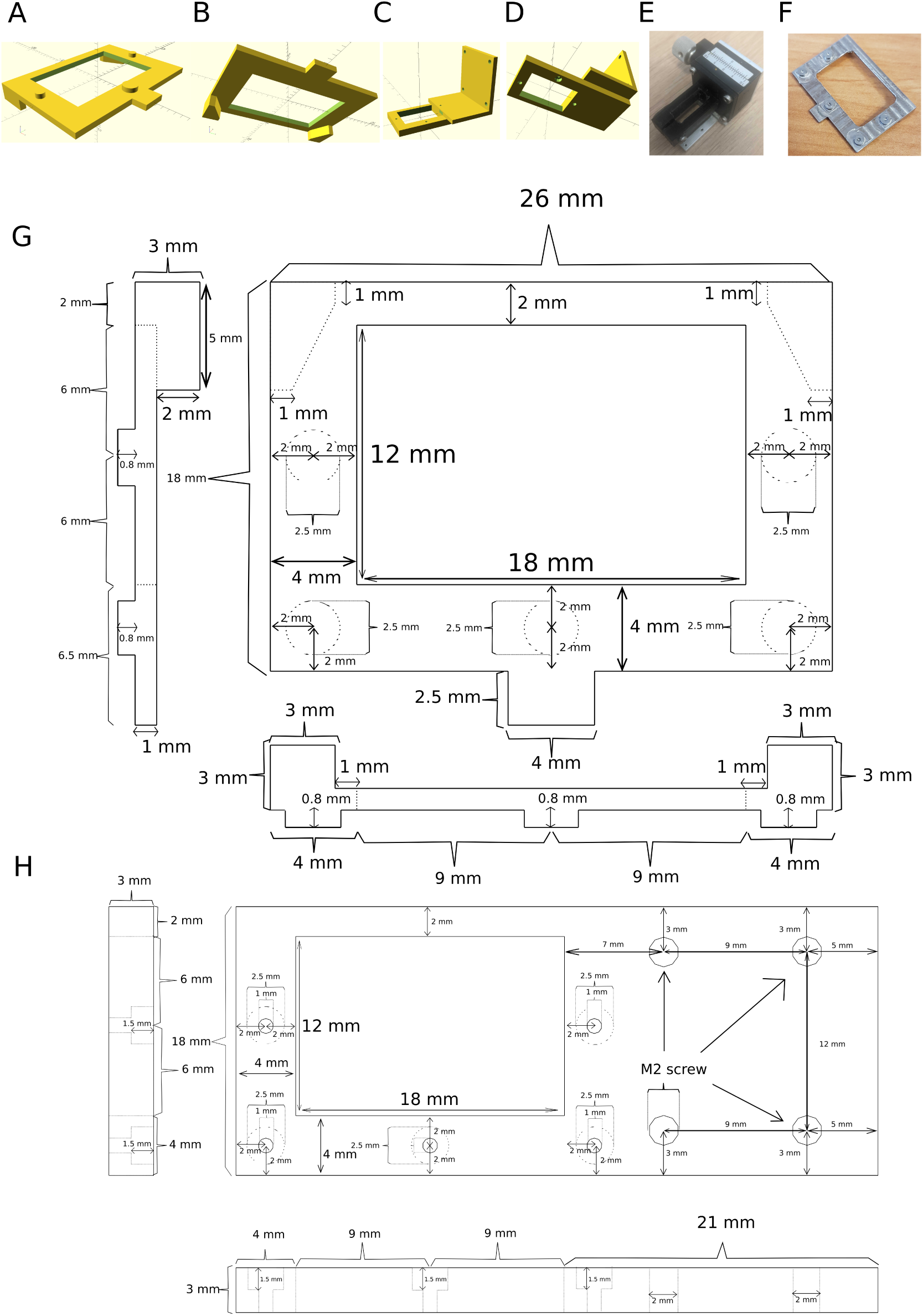
Design of the magnetic frame system for stabilizing the fUS recordings against movements. **A**: top view of the ferromagnetic frame. The frame was made of the corrosion resistant, ferromagnetic stainless steel 440C. **B**: bottom view of the ferromagnetic frame. The legs were used to aid in cementing by steadying the frame in place and providing locations to cement. **C**: top view of the magnetic frame. The frame was made of aluminum, and designed to hold the micromanipulator and supermagnets while otherwise minimizing the space taken up. **D**: Bottom view of the magnetic frame. The supermagnets were stored in the three holes on the bottom; a later version (blueprint in figure H) would have five holes for magnets. **E**: photo of magnetic frame with micromanipulator and the probe-clamp. **F**: photo of the 5-peg ferromagnetic frame. **G, H**: detailed blueprints for the ferromagnetic frame (G) and magnetic frame (H) given to the metal shop which constructed them.

**Supplementary Figure 2.**
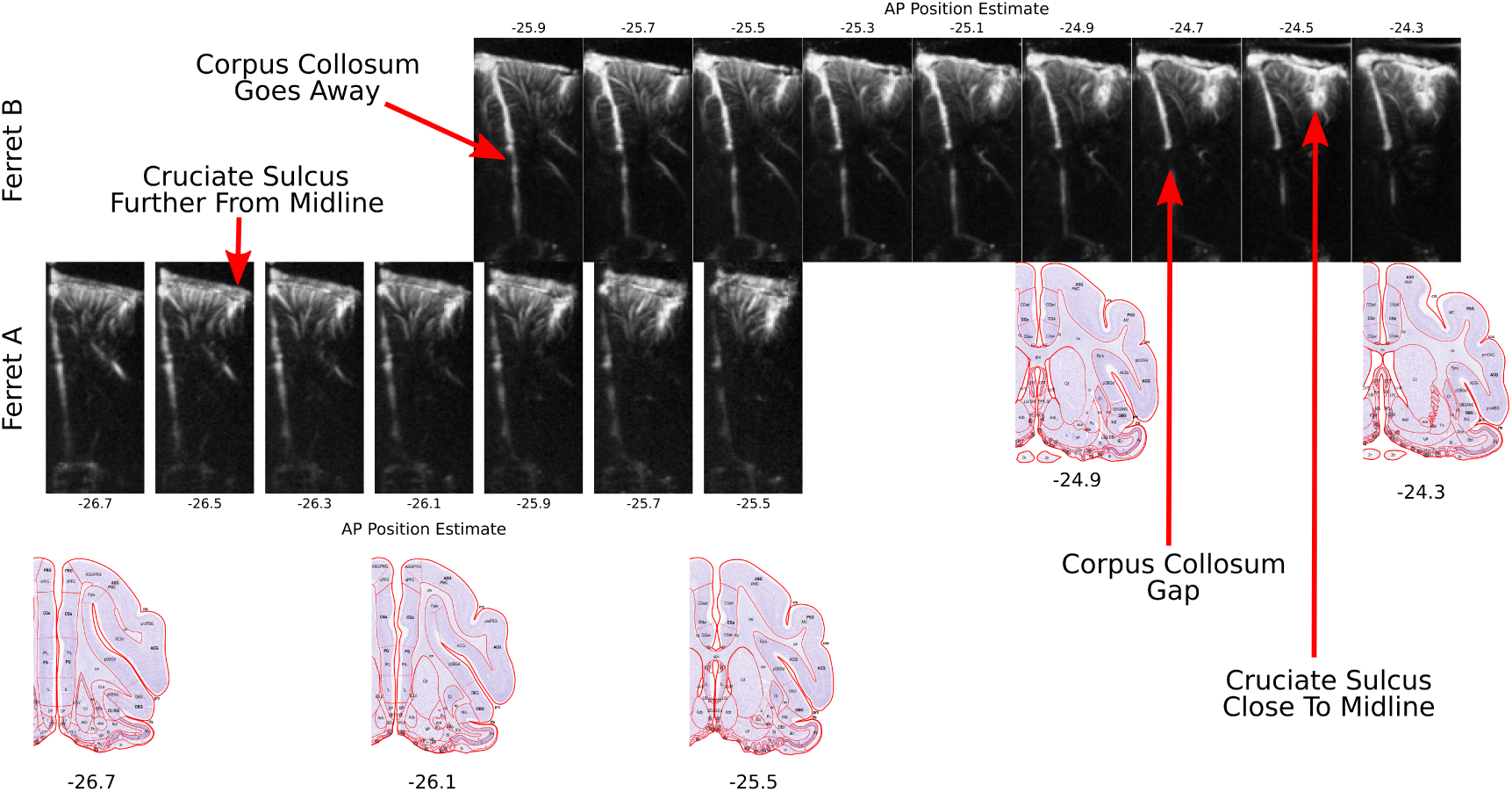
Demonstration of the method used for judging anatomical correspondences. The images for each ferret were taken at anterio-posterior steps of 200 micrometers. Images on the left are anterior, and on the right are posterior. In order to judge the anatomical location of any particular micromanipulator position, information from all anatomical images was taken into account. Particularly important landmarks in this case include the cruciate sulcus, which branches out from the interhemispheric fissure in a Y formation. Moving anterior from the branch point, it becomes more and more lateral. Also important and reliable as an anatomical anchor is the corpus collosum: when it is present, the large blood vessel in the intrahemispheric fissure has a “gap” in the image. Moving anterior, the corpus collosum is no longer present. This transition, according to the atlas, happens in between 26.1 and 25.5 mm anterior to the occipital crest. The example slice in the text was ferret A at -25.9.

**Supplementary Figure 3.**
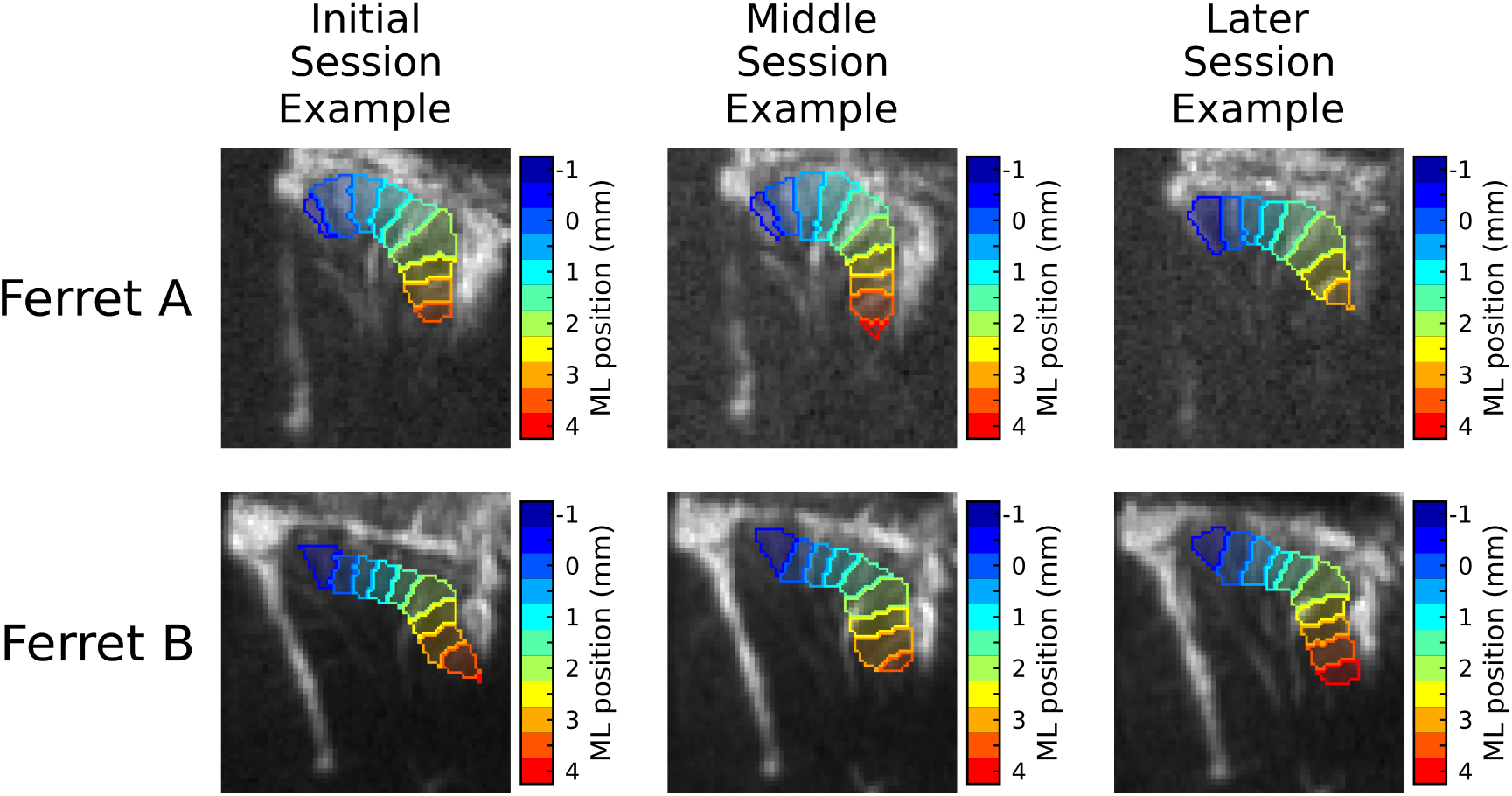
Consistency of anatomy and columns across recording sessions. Three example sessions are chosen for each ferret, using the same slice; the recordings were each taken approximately a week apart. Despite this, the gross anatomy looks the same for each image. Column positions were placed such that the positions of major vascular landmarks would be the same relative to each column; see for example the large vertical vessel under position "0" in ferret A, and the large vessel between columns 1.5 and 2.

**Supplementary Figure 4.**
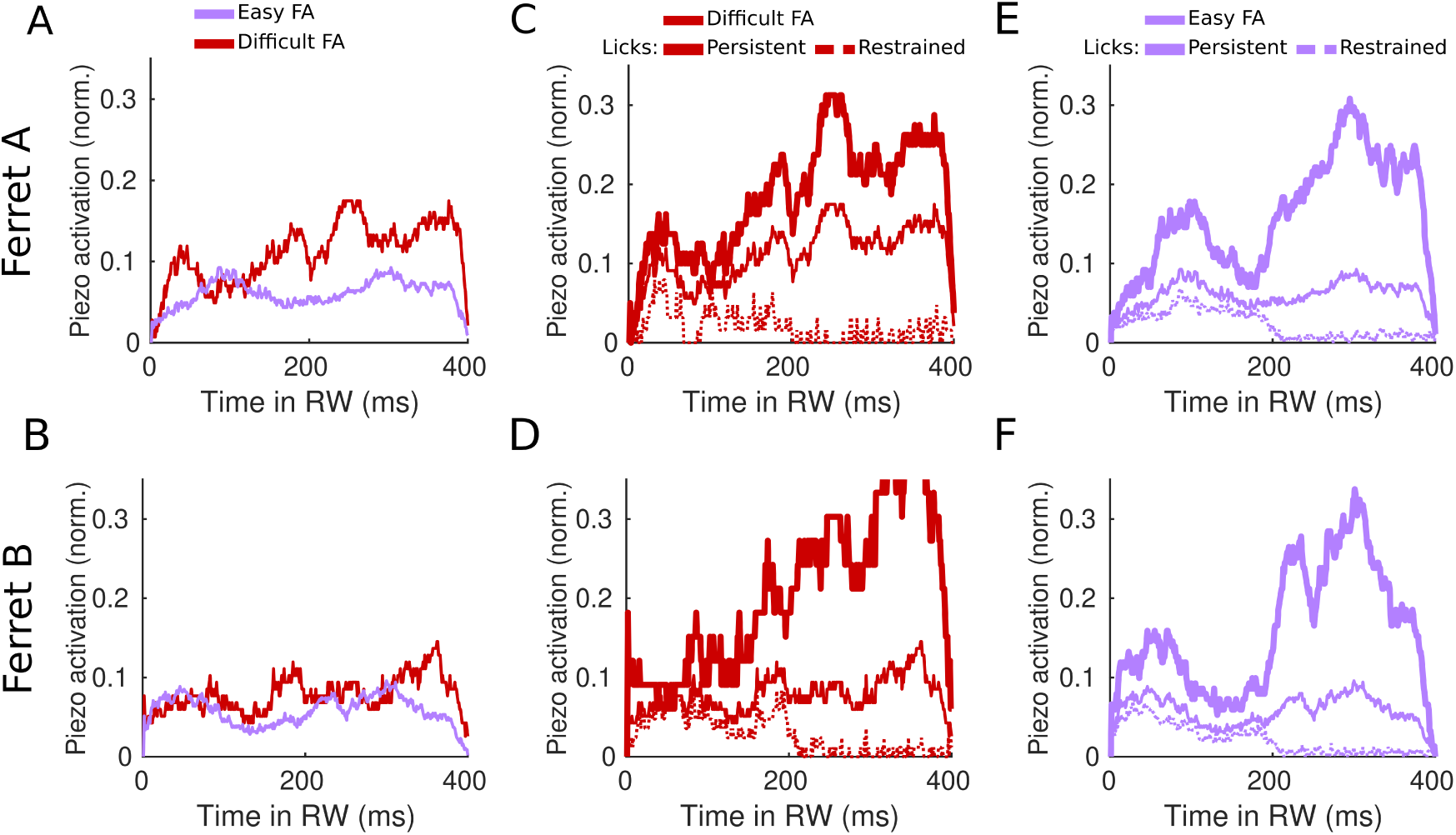
Illustration of the Persistent/Restrained Licking patterns after spout arrival. **A, B**: average lick rates during the response window for Easy and Difficult FA trials. **C, D**: Average lick rates for Persistent and Restrained Difficult FAs. **E, F**: Average lick rates for Persistent and Restrained Difficult FAs. Note that the rates of each category are approximately the same as the categories in panels C and D.

**Supplementary Figure 5.**
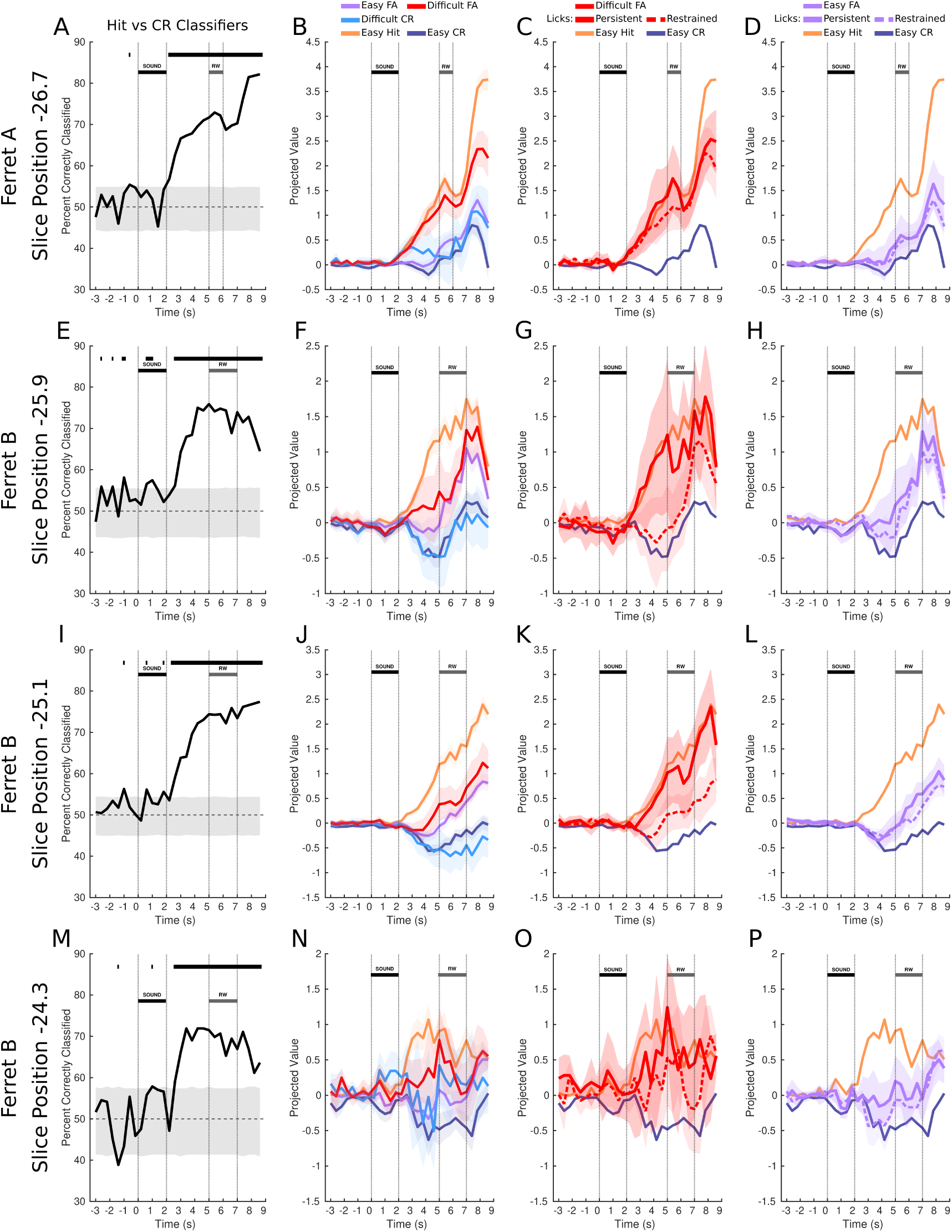
Replication of all example figures from Figures 1 2 and 3 for ferret A’s other slice and for ferret B’s slices. The ferret and estimated slice AP position are noted on the left of each row. **A, E, I, M**: percents correctly classified for each slice, as in figure 1I. **B, F, J, N**: projection plots as in Figure 2. Easy FA, Difficult FA, and Difficult CR projections are plotted on the same figure to save space. **C, G, K, O**: projection plots for Persistent/Restrained Lick trials among Difficult FA trials, as in Figure 3A. **D, H, L, P**: projection plots for Persistent/Restrained Lick trials among Easy FA trials, as in Figure 3B

**Supplementary Figure 6.**
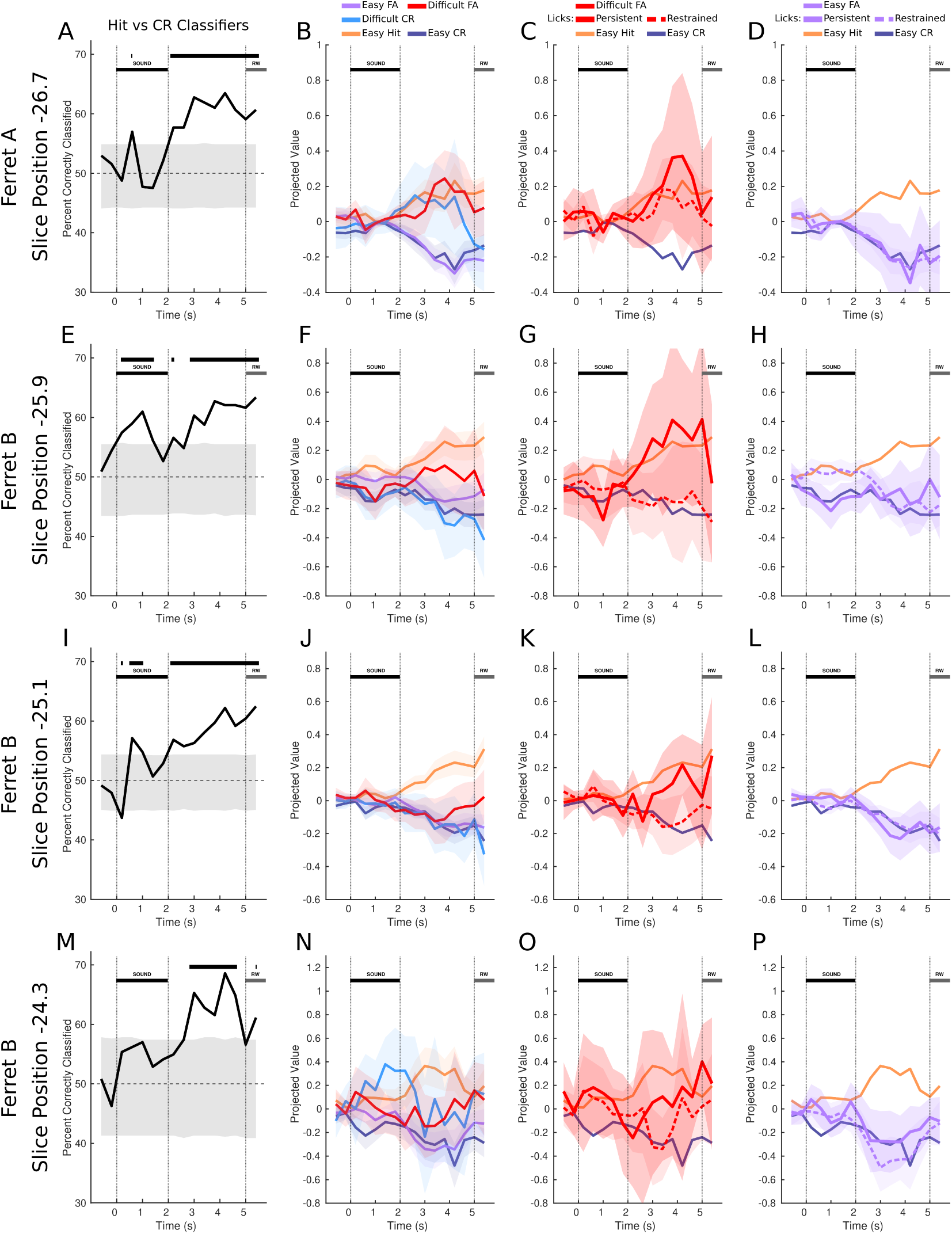
Same as Supplementary Figure 5, except after subtracting the regression model, as in Figure 4.

